# Identification of resistance mechanisms to small-molecule inhibition of TEAD-regulated transcription

**DOI:** 10.1101/2023.08.16.553491

**Authors:** Aishwarya Kulkarni, Varshini Mohan, Tracy T. Tang, Leonard Post, Murray Manning, Niko Thio, Benjamin L. Parker, Joseph Rosenbluh, Joseph H.A. Vissers, Kieran F. Harvey

**Author notes:** **Corresponding author:** Harvey, K.F.

## Abstract

The Hippo tumour suppressor pathway controls transcription by regulating nuclear abundance of YAP and TAZ, which activate transcription with the TEAD1-TEAD4 DNA-binding proteins. Recently, several small-molecule inhibitors of YAP and TEADs have been reported, with some now entering clinical trials for different cancers. Here, we investigated the cellular response to TEAD palmitoylation inhibitors, using a combination of genomic and genetic strategies. Genome-wide CRISPR/Cas9 screens identified genes that modulate the cellular response to TEAD inhibition, including members of the Hippo, MAPK and JAK-STAT signaling pathways. By exploring gene expression programs of mutant cells, we found that MAPK pathway hyperactivation confers resistance to TEAD inhibition by reinstating expression of a subset of YAP/TEAD target genes. Consistent with this, combined inhibition of TEAD and the MAPK protein MEK, synergistically blocked proliferation of several mesothelioma and lung cancer cell lines and more potently reduced the growth of patient-derived lung cancers in vivo. Collectively, we reveal mechanisms by which cells can overcome small-molecule inhibition of TEADs and potential strategies to enhance the anti-tumor activity of emerging Hippo pathway targeted therapies.

## INTRODUCTION

The Hippo pathway is an important regulator of organ growth and is also mutated in human cancers (Harvey *et al*, 2013). First discovered in *Drosophila*, it is highly conserved throughout evolution including in humans (Harvey & Hariharan, 2012; Zheng & Pan, 2019). The Hippo pathway has been predominantly studied in epithelial tissues, but also functions in other tissue types, including muscle, the nervous system and some blood cell types (Halder & Johnson, 2011). Early evidence of a role for the Hippo pathway in cancer came from studies whereby the human orthologue of Salvador (SAV1), a founding member of the *Drosophila* Hippo pathway, were found to be mutated in renal cancer cell lines (Tapon *et al*, 2002). More recently, large scale analyses of human cancer samples have revealed that multiple Hippo pathway genes are mutated in a broad range of cancer types (Wang *et al*, 2018). Whilst Hippo pathway gene perturbations are relatively rare in most cancer types, they are very common in select cancers, such as mesothelioma, meningioma and squamous epithelial cell cancers (Harvey *et al*., 2013; Kulkarni *et al*, 2020; Wang *et al*., 2018).

The Hippo pathway responds to many cell biological cues such as cell-cell adhesion, cell-extracellular matrix adhesion, cell polarity and mechanical cues (Davis & Tapon, 2019). It can also respond to various stresses such as changes in osmolarity, temperature and energy stress (Koo & Guan, 2018). The pathway is comprised of more than 40 proteins, that relay information from the plasma membrane to the nucleus in order to modulate transcription. Typically, these signals are transduced by a core kinase cassette consisting of the MST1/2 kinases (Hippo in *Drosophila*) and LATS1/2 kinases (Warts in *Drosophila*). The activity of these kinases is controlled by scaffold proteins such as SAV1, Mob1A/B (Mats in *Drosophila*) and Merlin (encoded by the neurofibromatosis type 2 gene *NF2*). These proteins control transcription by regulating the rate at which the YAP and TAZ (Yorkie in *Drosophila*) transcription co-activator proteins move between the nucleus and cytoplasm (Manning *et al*, 2020). YAP and TAZ regulate transcription by binding to the TEAD1-TEAD4 DNA binding proteins (Scalloped in *Drosophila*) and by recruiting transcriptional regulatory proteins such as the COMPASS complex, the Mediator complex and by promoting transcription elongation. TEAD1-4 can also repress transcription, together with the VGLL4 and INSM1 transcription repressor proteins (Zheng & Pan, 2019).

Given the prevalence of Hippo pathway deregulation in many cancers, the dramatic tissue overgrowth that is caused by mutation of Hippo pathway genes in model organisms like *Drosophila* and mice, and the paucity of effective therapies for cancers like mesothelioma, the Hippo pathway is considered a potential therapeutic target. Interestingly, unbiased screening efforts by both pharma and academia have identified TEAD1-4 as druggable Hippo pathway proteins (Calses *et al*, 2019; Dey *et al*, 2020; Pobbati *et al*, 2023). Many compounds that influence TEAD-regulated transcription work by modulating auto-acylation of a hydrophobic pocket in TEADs that normally stabilizes them and promotes their physical association with YAP and TAZ (Chan *et al*, 2016; Holden *et al*, 2020). Multiple high potency TEAD inhibitors (TEADi) have now been developed and subsequently entered clinical trials for cancers such as mesothelioma, as well as other cancers that harbor *NF2* mutations or oncogenic YAP gene fusions (Calses *et al*., 2019; Dey *et al*., 2020; Pobbati *et al*., 2023). However, given the relatively recent discovery of TEADi, we lack a detailed understanding of their mechanism of action. In addition, although targeted therapies have revolutionized cancer treatment, therapy resistance is inevitable in almost all patients and thus limits their long-term effectiveness. As such, although Hippo-targeted therapies are predicted to be powerful anti-cancer agents, some degree of resistance to them is likely. To explore mechanisms of cell-intrinsic resistance to TEADi, and thus identify potential therapies that can be used in combination with TEADi, we performed whole genome CRISPR/Cas9 screens in mesothelioma cells. Multiple genes were identified that modulate the cellular response to TEAD inhibitors, most notably members of the Hippo, MAPK and JAK/STAT pathways, thus identifying potential combination therapy approaches to be coupled with TEADi.

## RESULTS

The discovery that TEAD transcription factors are post-translationally modified via auto-palmitoylation, and the importance of this for YAP/TAZ-TEAD driven transcription (Chan *et al*., 2016; Noland *et al*, 2016), has aided the development of small-molecule inhibitors of TEADs. The majority of these inhibitors block auto-palmitoylation of key cysteine residues in TEADs, impact the ability of TEADs to bind to YAP, and/or block proliferation of cancer cells with Hippo pathway mutations (Bum-Erdene *et al*, 2019; Chan *et al*., 2016; Gibault *et al*, 2021; Gridnev *et al*, 2022; Holden *et al*., 2020; Hu *et al*, 2022; Kaneda *et al*, 2020; Laraba *et al*, 2023; Li *et al*, 2020; Lu *et al*, 2021; Lu *et al*, 2023; Lu *et al*, 2019; Noland *et al*., 2016; Pobbati *et al*, 2015; Sun *et al*, 2022; Tang *et al*, 2021). Here, we used a range of approaches including large-scale functional genomics screens and transcriptome profiling to investigate the mechanism of action of a potent small molecule inhibitor of all four TEADs (VT107), developed by Vivace Therapeutics (Tang *et al*., 2021). After confirming the nanomolar-potency of VT107 in the Hippo-pathway mutant mesothelioma cell lines NCI-H2052 (*NF2*, *LATS2* mutant) and NCI-H226 (*NF2* ^-/-^) (Miyanaga *et al*, 2015; Murakami *et al*, 2011; Sekido *et al*, 1995) (Fig. 1A-B), we used proteomics analyses and RNA sequencing (RNA-seq) to validate its on-target activity (Fig. 1C-H, Fig. S1, Table S1, S2). Gene expression was strongly impacted in both cell lines by VT107 and many known YAP/TAZ-TEAD target genes (e.g., *NPPB, IGFBP3, SNAPC1, CTGF*, *CYR61*, and *ANKRD1)* were among the top downregulated genes (Fig. 1D-E) (Cordenonsi *et al*, 2011; Hao *et al*, 2008; Varelas *et al*, 2008; Zanconato *et al*, 2015; Zhao *et al*, 2008). This was further confirmed by Gene set enrichment analysis (GSEA), which indicated that YAP activity was strongly reduced by VT107 in both cell-lines (Fig. 1F) (Cordenonsi *et al*., 2011; Subramanian *et al*, 2005). Finally, immunoblotting and proteomics experiments revealed that the expression of YAP/TAZ targets was reduced at the protein level by VT107 (Fig. 1G-H, Fig. S1, Table S2). In addition to limiting the expression of YAP-TEAD target genes, VT107 modulated the expression of many genes that are not known to be regulated by YAP-TEAD. Most notably, genes that are responsive to RAS/MAPK pathway activity and cytokine signaling (e.g., IL2 and IL15) were strongly upregulated upon VT107 treatment, while genes linked to the serum response, Hedgehog pathway and ribosome biogenesis were downregulated (Fig. 1F).

**Figure 1.**
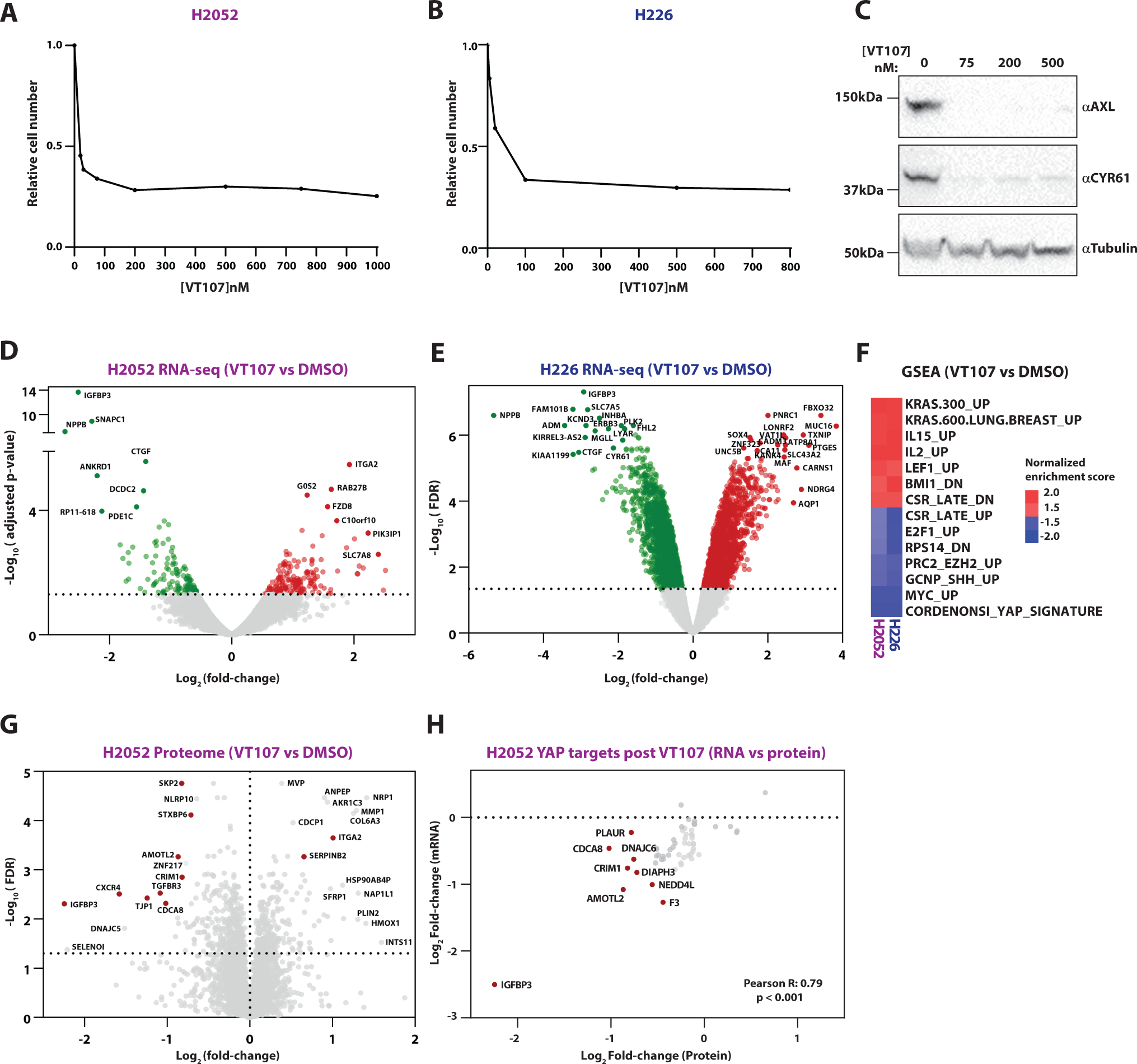
The effect of TEAD inhibitors on the transcriptome and proteome of Hippo pathway mutant mesothelioma cells. **A-B)** A chart of H2052 (A) and H226 (B) cell numbers following treatment with different doses of VT107 (between 5-2500nM) for 6 days under CRISPR/Cas9 screening conditions, n=1. **C)** Immunoblots of lysates from H2052 cells treated with different doses of VT107 for 6 days and probed with the indicated antibodies. **D-E)** Volcano plots of gene expression of H2052 and H226 cells following 24-hour VT107 treatment, n=3. Genes whose expression was strongly modulated are labelled. **F)** A heatmap of strongly modulated transcriptional signatures by VT107 in H2052 and H226 cells, as determined by gene set enrichment analysis of differentially expressed genes (VT107 vs DMSO). **G)** A volcano plot of protein expression of H2052 cells following 24-hour VT107 treatment, n=5. YAP/TAZ-TEAD targets are highlighted in red. **H)** A correlation analysis of the effect of VT107 on mRNA and protein abundance of YAP/TAZ-TEAD targets. All plotted targets have significant protein level changes (q-value ≤ 0.05) in response to VT107. Correlation was assessed using the Pearson’s correlation coefficient. Most significantly changed targets are labeled.

### Unbiased identification of genes that modulate the cellular response to TEAD inhibitors

To identify genes that either enhance or suppress the cellular response to VT107, we performed genome-wide CRISPR/Cas9 screens in both NCI-H2052 (H2052) and NCI-H226 (H226) cells (Fig. 2A). Each screen was conducted in duplicate using the Brunello CRISPR library and two doses of VT107 applied sequentially for up to two weeks per dose (Doench *et al*, 2016; Doench *et al*, 2014). Cells were treated first with an IC_50_ dose (18nM in H2052 cells, 32nM in H226) and then with a cytostatic dose (100nM for each cell line). Whole-genome sequencing was performed on genomic DNA extracted at the endpoint of vehicle (DMSO) or VT107 treatment (at both concentrations) and sgRNA abundance was quantified. Using the MAGeCK algorithm we first compared sgRNA abundance in DMSO treated cells and the initial DNA pool (Li *et al*, 2014). The abundance of sgRNAs targeting known core cell essential genes (Hart *et al*, 2015) was decreased at the screening endpoint (Fig. S2A-B, Table S3), and we observed strong correlations in the gene log fold-changes in DMSO-treated populations with CRISPR/Cas9 screens of H2052 and H226 cells from the DepMap database, indicating the technical success of our screen (Fig. S2C-D, Table S4).

**Figure 2.**
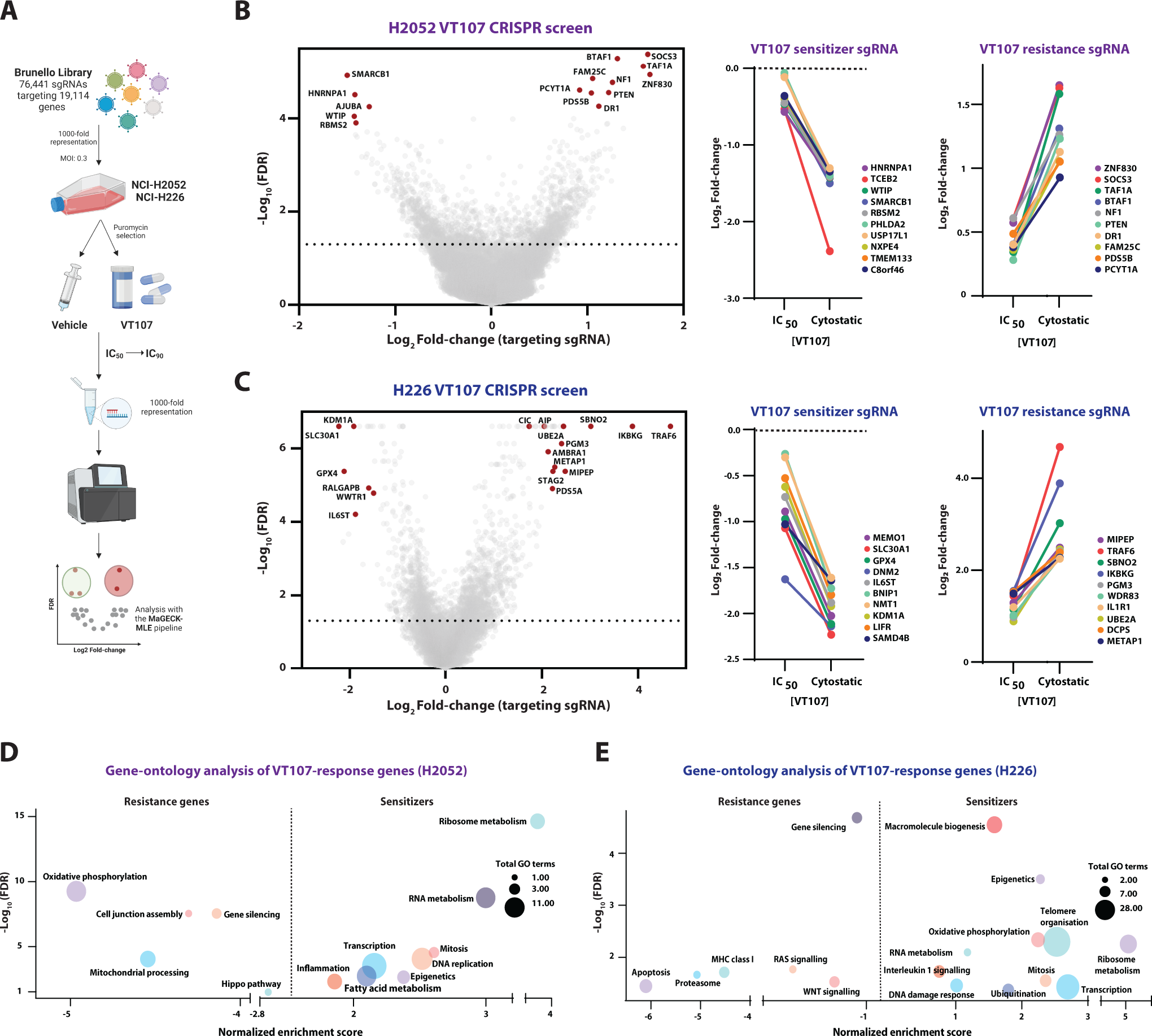
Identification of genes that modulate the response of mesothelioma cells to TEAD inhibitors. **A)** A schematic diagram outlining the protocol for the VT107 CRISPR/Cas9 screens. **B-C)** Left panels: volcano plots representing the depletion or enrichment of targeting sgRNA (per gene) following treatment with 100nM VT107 (cytostatic dose) in CRISPR/Cas9 screens in H2052 cells (B) and H226 cells (C). Comparison of the depletion (middle panels) or enrichment (right panels) of targeting sgRNA following treatment with VT107 at IC_50_ and cytostatic doses in H2052 cells (B) or H226 cells (C) in CRISPR/Cas9 screens. **D-E)** Bubble charts representing ontology analysis categories of genes that conferred VT107 resistance (Targeting sgRNA log_2_Fold-change < 0) or sensitivity (Targeting sgRNA log_2_Fold-change > 0) in CRISPR/Cas9 screens in H2052 cells (D) or H226 cells (E). Each bubble is plotted according to the strongest scoring GO term from its respective category and total GO terms per category are indicated by bubble size.

To identify genes that confer resistance to or synergize with VT107 after CRISPR mediated gene disruption, we used the MAGeCK algorithm to compare sgRNA abundance in DMSO treated cells and VT107-treated cells. We filtered genes whose targeting-sgRNA showed significant (FDR ≤ 0.05) changes in abundance at both screening endpoints and applied a magnitude threshold to identify genes whose loss either conferred resistance to VT107 (targeting-sgRNA log fold-change ≥ 0.5 in VT107-treatment population relative to vehicle population) or sensitivity to VT107 (targeting-sgRNA log fold-change ≤ -0.5). This revealed 674 or 818 putative drivers of VT107 sensitivity and 571 or 519 VT107 resistance genes in H2052 and H226 cells, respectively (Fig. 2B-C, Table S5). Guide RNAs targeting the majority of VT107 sensitivity genes showed a positive fold-change at both the IC_50_ and IC_90_ timepoints, and similarly, gRNAs targeting most VT107 resistance genes exhibited negative log fold-changes at both doses (middle and left panels of Fig. 2B-C, and Fig. S2E-H). This indicated that the response to VT107 was maintained for the duration of both screens. Putative sensitivity genes from both screens were enriched in the cellular processes of ribosome metabolism, RNA metabolism and transcription, whilst some ontology groups were only enriched in one cell line (Fig. 2D-E).

By comparing the H2052 and H226 CRISPR/Cas9 screens, we found a weak positive correlation (Pearson’s R: 0.1921, p < 0.01), further indicating shared mechanisms in how these cells respond to VT107, but also differences (Fig. 3A). Common sensitivity and resistance genes from each screen were identified, as well as common gene ontology groups (Fig. 3A-C). Among these, several key Hippo pathway genes modulated the response to VT107, most notably *VGLL4* (whose loss promoted VT107 resistance) and *WWTR1 (*the loss of which promoted VT107 sensitivity) (Fig. 3A-B). *VGLL4* encodes a transcriptional repressor protein that competes with YAP for TEAD1-4 binding, whilst *WWTR1 (*also known as *TAZ)* is a paralog of *YAP* (Hong *et al*, 2005; Koontz *et al*, 2013). Upstream Hippo pathway genes that promote pathway activity were not identified as conferring resistance to VT107, possibly because the Hippo pathway is already strongly perturbed in H2052 and H226 cells, owing to *NF2* and *LATS2* mutations and/or because of functional redundancy between homologous genes (Miyanaga *et al*., 2015; Murakami *et al*., 2011; Sekido *et al*., 1995). In fact, the loss of some upstream Hippo pathway genes (e.g., *PTPN14*) counterintuitively conferred some degree of sensitivity to VT107 (Fig. 3B), the significance of which is currently unclear (Wang *et al*, 2012). In addition, sgRNA-based depletion of the LATS kinase repressors WTIP, AJUBA and TRIP6 conferred sensitivity to VT107 in H2052 but not H226 cells (Fig. 2B, Table S5). Strikingly, two major negative regulators of the MAPK pathway and bona fide human tumor suppressor genes, *Neurofibromin 1* (*NF1*) and *Capicua* (*CIC*), conferred strong resistance to VT107 when depleted by sgRNA in both cell lines (Fig. 3A-B) (Cichowski & Jacks, 2001; Kawamura-Saito *et al*, 2006; Kim *et al*, 2021; LeBlanc *et al*, 2017; Ratner & Miller, 2015; Xu *et al*, 1990). In addition, inactivation of the JAK-STAT pathway effector *STAT3* conferred strong sensitivity to VT107, while that of the JAK-STAT pathway repressor *SOCS3* elicited strong resistance in H2052 cells (Fig. 3A-B) (Rawlings *et al*, 2004). Collectively, genome-wide CRISPR/Cas9 screens indicated that modulation of multiple genes and signaling pathways can modify the response of mesothelioma cells to small molecule inhibition of YAP/TEAD-mediated transcription.

**Figure 3.**
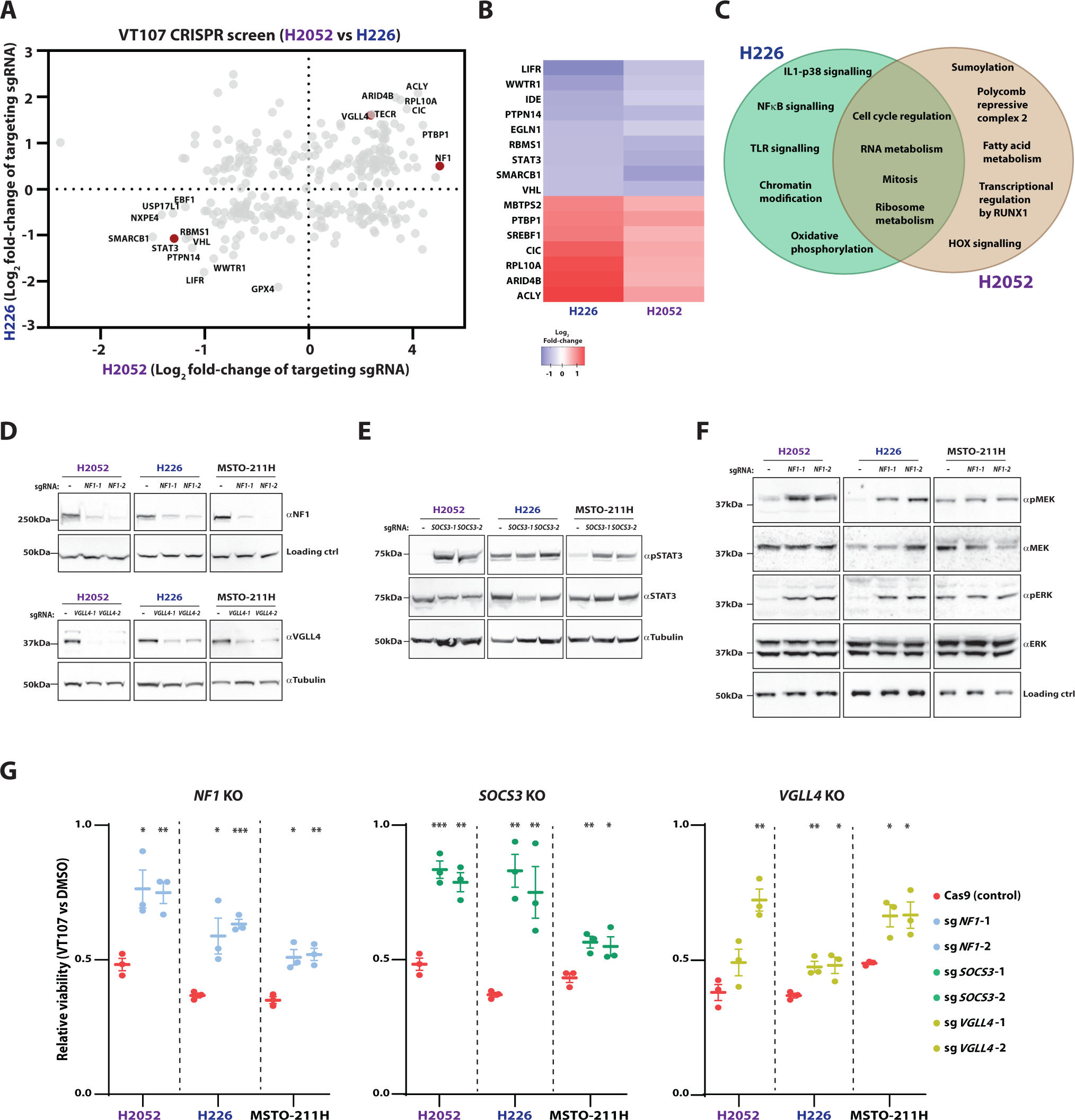
Inactivation of MAPK, JAK/STAT and Hippo pathway repressors confers resistance to TEAD inhibitors. **A)** A correlation plot comparing the response to VT107 treatment in CRISPR/Cas9 screens in H2052 and H226 cells. Genes that are highlighted in red were subsequently studied. Correlation was assessed using the Pearson’s correlation coefficient. **B)** A heatmap of high stringency VT107 resistance genes (Targeting sgRNA log_2_Fold-change ≤ -0.8) and sensitivity genes (Targeting sgRNA log_2_Fold-change ≥ 0.8) identified in CRISPR/Cas9 screens in both H2052 and H226 cells. **C)** A venn diagram of GO terms that were enriched among VT107 resistance or sensitivity genes in CRISPR/Cas9 screens in either H2052 or H226 cells alone, or common to both screens. **D-F)** Immunoblots of lysates from H2052, H226 and MSTO-211H cells expressing Cas9. Cells were control parental cells or expressed the following independent gRNAs: *NF1* in D (top panel) and F; *VGLL4* in D (bottom panel); *SOCS3* in E. Lysates were probed with the indicated antibodies, while detection of a non-specific protein serves as a loading control in D (top panel) and in F. Molecular mass markers are indicated. **G)** Charts of the impact of VT107 on the viability of parental and mutant H2052, H226 and MSTO-211H cells, as assessed by alamar blue assays. H2052 and H226 cells were treated with 100nM VT107 and MSTO-211H cells with 370nM VT107. Genes that were mutated were *NF1* (left panel), *SOCS3* (middle panel) or *VGLL4* (right panel). n=3, error bars indicate SEM, student’s t-tests were used for all statistical comparisons between parental and mutant cells, * p < 0.05, ** p < 0.01, *** p <0.001.

### Inactivation of MAPK, JAK/STAT and Hippo pathway repressors confers resistance to TEAD inhibitors

To validate our whole-genome screens, we used CRISPR/Cas9 mutagenesis with two independent sgRNAs to mutate *NF1*, *SOCS3* and *VGLL4* in H226 and H2052 cells, as well as a third mesothelioma cell line, MSTO-211H, which harbors inactivating mutations in both *LATS1* and *LATS2* (Miyanaga *et al*., 2015). A substantial reduction of NF1 and VGLL4 protein was confirmed by western blot in all 6 mutant cell lines (Fig. 3D). Endogenous SOCS3 protein could not be detected by western blot so we used Tracking of Indels by Decomposition (TIDE) analysis to confirm CRISPR/Cas9 mutagenesis at the *SOCS3* locus in all sg*SOCS3*-transduced cell lines (Fig. S3). Additionally, we assessed JAK/STAT pathway activation using STAT3 and p-STAT3 antibodies (STAT3 becomes phosphorylated on Y705 when the pathway is activated) (Heinrich *et al*, 1998) and observed strong JAK/STAT pathway activation in *SOCS3* mutant H2052 and MSTO-211H cells, while pathway activity was already elevated in H226 cells (Fig. 3E). *NF1* loss caused strong MAPK pathway hyperactivation in both H2052 and H226 cells (as assessed by phosphorylation of MEK and ERK at S221 or T202/Y204, respectively), while MSTO-211H cells exhibited high baseline MAPK pathway activity (Fig. 3F). Importantly, mutation of *NF1* or *SOCS3* with both sgRNAs induced VT107 resistance in all three cell lines, while mutation of *VGLL4* with one sgRNA conferred resistance to VT107 in all three cell lines, and in 2 out of 3 cell lines with the other sgRNA (Fig. 3G).

### Mutation of *NF1* reinstates expression of select YAP/TAZ-TEAD target genes

Next, we sought to investigate how resistance genes identified in our CRISPR/Cas9 screens modulate the cellular response to TEAD inhibition. Given the close functional links between the MAPK and Hippo pathways in transcription and cancer (Herranz *et al*, 2012; Koo *et al*, 2020; Lin *et al*, 2015; Pascual *et al*, 2017; Pham *et al*, 2021; Reddy & Irvine, 2013; Stein *et al*, 2015; Zanconato *et al*., 2015), we investigated this in *NF1* mutant H2052 cells, which displayed strong MAPK pathway activation (Fig. 3F). Parental and *NF1* mutant H2052 cells were treated with DMSO or VT107, and RNA harvested and sequenced 24 hours later. As expected, the transcriptome of *NF1* mutant cells differed significantly from parental cells in the baseline (DMSO-treated) condition (Fig. 4A and S4A). Unbiased gene set enrichment analyses revealed elevation of transcriptional signatures linked to the WNT and Estrogen response pathways in *NF1* mutant cells, consistent with published reports (Luscan *et al*, 2014; Zheng *et al*, 2020). Transcription signatures associated with oxidative phosphorylation and fatty acid metabolism were also elevated in *NF1* mutant cells (Fig. 4B).

**Figure 4.**
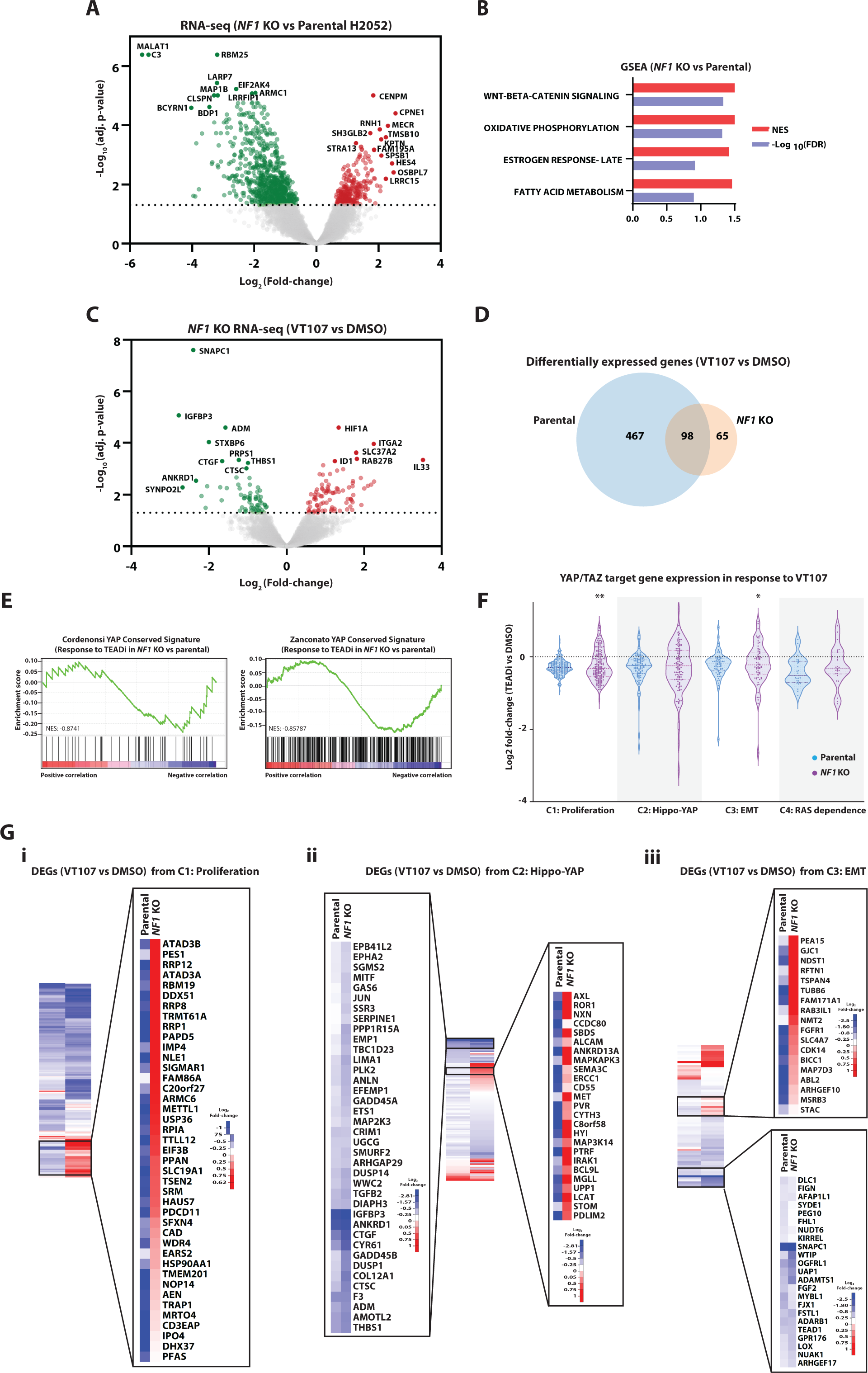
Mutation of *NF1* in mesothelioma cells reinstates expression of select YAP/TAZ-TEAD target genes. **A)** A volcano plot of gene expression of *NF1* mutant cells relative to parental H2052 cells, n=3. **B)** A bar chart indicating top scoring transcription signatures from gene set enrichment analysis of differentially expressed genes in *NF1* mutant cells compared to parental cells. **C)** A volcano plot of gene expression of *NF1* mutant H2052 cells (treated with DMSO or 1µM VT107 for 24 hours), n=3. **D)** A Venn diagram indicating the degree of overlap of differentially expressed genes in parental and *NF1* mutant cells treated with VT107. **E)** Gene set enrichment analysis plots of two YAP transcription signatures among differentially expressed genes in VT107-treated *NF1* mutant compared to parental H2052 cells. **F)** Violin plots of differentially expressed genes (VT107 vs DMSO) in both parental and *NF1* mutant H2052 cells. Four independent clusters of genes that are sensitive to YAP/TAZ expression were assessed. The null hypothesis for differences between parental and *NF1* mutant cells was tested with a hypergeometric distribution analysis ** p < 0.01, * p < 0.05. **E)** Heatmaps of relative gene expression level changes (VT107 vs DMSO) in both parental and *NF1* mutant H2052 cells. Heatmaps are shown for genes from cluster 1 (i), cluster 2 (iii) and cluster 3 (iii). Magnified regions focus on select genes from clusters 1-3 whose expression was restored in VT107-treated *NF1* mutant, compared to parental, H2052 cells. Also magnified are select genes from clusters 2 and 3 whose expression was not restored by *NF1* loss; these include many core YAP/TAZ-TEAD target and Hippo pathway genes.

Next, we compared the transcriptional response to VT107 in parental and *NF1* mutant H2052 cells. VT107 had a more profound effect on the transcriptome of parental cells compared to *NF1* mutant cells; expression of 564 genes was significantly altered by 24h of VT107 treatment in parental cells whilst only 163 genes were altered by VT107 in *NF1* mutant cells (Fig. 1D, 4C and Table S6). Despite the change in magnitude of gene expression changes following VT107 treatment between *NF1* mutant and parental cells, the changes were qualitatively similar: 98 genes displayed similar expression changes, including both downregulated (*IGFBP3*, *SNAPC1*, *ANKRD1*) and upregulated (*ITAG2*) genes (Fig. 1D, 4C-D and Table S6).

Given that the Hippo and MAPK pathways co-regulate many genes (Koo *et al*., 2020; Obier *et al*, 2016; Pascual *et al*., 2017; Pham *et al*., 2021; Stein *et al*., 2015; Zanconato *et al*., 2015), and VT107’s impact on the transcriptome was blunted in *NF1* mutant H2052 cells, we considered the possibility that expression of YAP-TEAD target genes was restored in VT107-treated *NF1* mutant cells. To investigate this, we performed targeted GSEA with published YAP-TEAD target gene signatures (Cordenonsi *et al*., 2011; Pham *et al*., 2021; Zanconato *et al*., 2015). Subsets of genes from two independent YAP-TEAD signatures were depleted less in VT107-treated *NF1* mutant cells relative to parental cells, implying weakly restored expression of YAP-TEAD target genes (Fig. 4E). To examine this further, we assessed the expression of four published clusters of genes that were recently reported to be strongly associated with depletion of YAP/TAZ across multiple cell types (Pham *et al*., 2021). Functionally, these clusters are most closely related to: 1) cell proliferation, including genes regulated by the Myc oncoprotein; 2) Hippo signaling (Hippo-YAP); 3) epithelial to mesenchymal transition (EMT); and 4) MAPK signaling (RAS dependence) (Pham *et al*., 2021). By comparing the magnitude of expression changes of each gene in all four clusters upon VT107 treatment, we found that select genes from each cluster were restored in *NF1* mutant cells (Fig. 4F). Statistical analyses revealed that cluster 1 genes and cluster 3 genes, when considered as populations, were significantly restored in *NF1* mutant cells treated with VT107, compared to parental cells (Fig. 4G). Finally, we examined expression changes in individual genes from select clusters, which underscored the observation that only select genes that are responsive to YAP/TAZ-regulated transcription were restored in *NF1* mutant cells upon VT107 treatment (Fig. 4G). Interestingly, the expression of many genes from cluster 2 that are considered “core” YAP/TAZ-TEAD target genes and are often used as surrogates for YAP activity (e.g., *CTGF, CYR61, ANKRD1, AMOTL2*) were not restored in *NF1* mutant cells, while many other genes were (Fig. 4G). Similarly, many genes that encode Hippo pathway proteins from the EMT cluster (e.g., TEAD1, WTIP, FJX1, ARHGEF17 and KIRREL) were not restored in VT107-treated *NF1* mutant H2052 cells. Collectively, these studies suggest that VT107 resistance in *NF1* mutant cells was driven by partial reinstatement of genes that are sensitive to YAP/TAZ activity.

### Combined MAPK and Hippo pathway inhibition synergistically impacts mesothelioma cells

Next, we sought to determine whether combining MAPK pathway or JAK/STAT pathway inhibition with VT107 enhanced its ability to inhibit mesothelioma cell proliferation. For this, we measured the impact of combining VT107 treatment with either the MEK1/2 inhibitor trametinib or 5 independent JAK-STAT inhibitors that target different components of the IL6/Gp130-JAK-STAT3 pathway (Fig. 5A). In both H2052 and H226 cell lines, VT107 and trametinib synergistically inhibited cell proliferation (Fig. 5B-D), which is consistent with several published studies on genetic or chemical targeting of the Hippo and MAPK pathways (Kim *et al*, 2016; Koo *et al*., 2020; Lin *et al*., 2015; Park *et al*, 2020; Zanconato *et al*., 2015). In contrast, while the JAK1/2 inhibitor AZD1480 exhibited synergistic or strong additive interactions with VT107 in H226 and H2052 cells (Fig. 5B, 5E-F), the other JAK-STAT pathway inhibitors we tested exhibited weak additive interactions when combined with VT107 in both cell lines (Fig. 5B and Fig. S5). Next, we investigated how general the impact of combining Hippo/MAPK therapies is by expanding our studies to include additional mesothelioma cell lines and an independent but related TEADi, VT108. VT108 and trametinib synergistically impeded proliferation of four out of 10 mesothelioma cell lines tested and had additive impacts on the proliferation of the remaining 6 cell lines (Fig. 5G and Fig. S5Bi).

**Figure 5.**
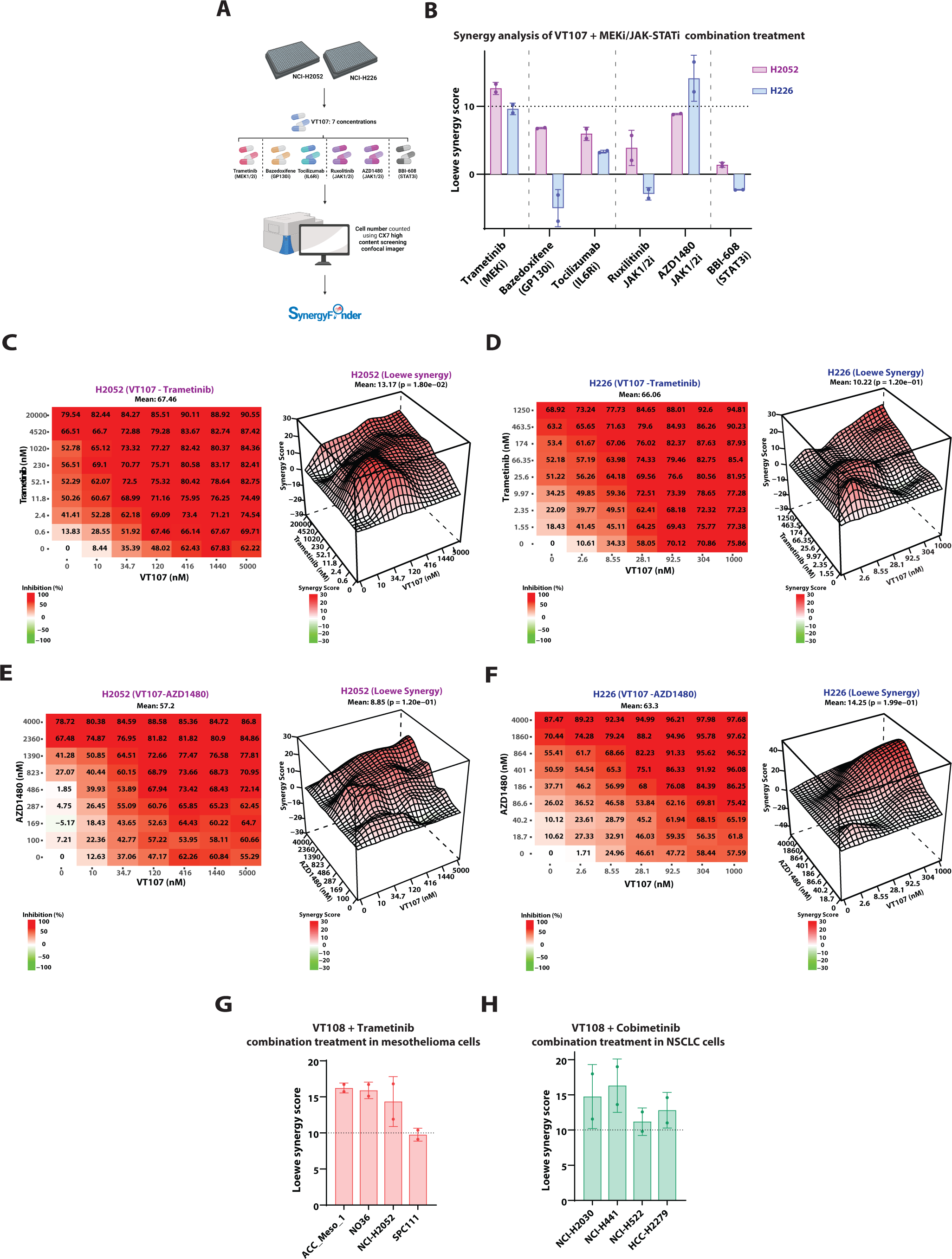
Combined MAPK and Hippo pathway inhibition synergistically impacts mesothelioma and NSCLC cells. **A)** A schematic diagram outlining the protocol used for combination drug treatment assays conducted in H2052 or H226 cells. **B)** A bar chart indicating the interaction outcome of combined treatment with VT107 and the indicated compounds on H2052 and H226 cell numbers. Synergistic interactions were indicated by synergy scores ≥ 10 and additive interactions by scores within the range of -10 to 10. n=2 biological replicates (2 technical replicates per biological replicate), error bars: SEM. **C-F)** Left panels: dose response matrices indicating the level of inhibition of cell number by treatment of cells with the indicated doses of VT107 and/or Trametinib. Right panels: Representative 3D synergy calculation topological maps indicating the type and degree of interaction between drugs on inhibition of cell number. H2052 cells were treated in C and E, H226 cells were treated in D and F. VT107 was combined with MEKi in C and D, and with AZD1480 in E and F. Mean synergy scores were assessed statistically using parametric bootstrapping method, n=2. **G-H)** Bar charts indicating the interaction outcome of combination treatment with Left: VT108 and Trametinib in mesothelioma cells or Right: VT108 and Cobimetinib in NSCLC cells. In all assays, n=2 technical replicates and error bars: SD.

### Combined MAPK and Hippo pathway inhibition synergistically impacts non-small cell lung cancer cells and tumors

To explore combined inhibition of the Hippo and MAPK pathways in non-mesothelioma cancers, we performed similar combination therapy studies on non-small cell lung cancer (NSCLC) cell lines. In addition, to determine whether the synergistic impact of combined Hippo/MAPK therapy was not limited to the MEKi trametinib, we utilized an independent MEKi, cobimetinib (Rice *et al*, 2012). Synergistic impacts between VT108 and cobimetinib were observed in ten out of 42 NSCLC cell lines tested, while additive effects were observed on the proliferation of the remaining 32 cell lines (Fig. 5H and Fig. S5B). This indicates that TEADi and MEKi can also synergistically impact cancer cells of non-mesothelioma origin.

Given the strong support for combined Hippo and MAPK pathway inhibition as a cancer treatment in our studies and in the literature, we explored the effect of combined TEADi and MEKi treatment on cancer cell line-derived tumors grown *in vivo*. For this, we selected two patient-derived xenograft (PDX) models of *NF2* mutant NSCLC cells, as we observed TEADi-MEKi synergism in both NSCLC and mesothelioma cell lines *in vitro* (Fig. 5, S5B), and Hippo mutant mesothelioma cells either did not form tumors in mice or were unsuitable for in vivo combination therapy experiments as they were extremely sensitive to TEADi monotherapy (Tang *et al*., 2021). PDX tumor-bearing mice were treated with either vehicle, trametinib alone, VT108 alone, or a combination of both compounds, with doses defined based on preliminary experiments with each tumor type. As monotherapies, trametinib and VT108 reduced tumor growth to some extent in both PDX tumors (Fig. 6A, 6C). For LU-01-0407 PDX tumors, combination trametinib and VT108 therapy reduced tumor size significantly more than either agent alone or led to almost complete tumor regression (Fig. 6A). For LU-01-0236 PDX tumors, combination trametinib and VT108 therapy reduced tumor growth significantly more than VT108 alone (Fig. 6C). In addition, we observed no unfavorable impact on the body weight of mice throughout the course of the experiment (Fig. 6B, 6D). Collectively, these tumor xenograft studies provide in vivo evidence for the benefit of treating Hippo mutant tumors with combination therapies that target both MEK and TEAD.

**Figure 6.**
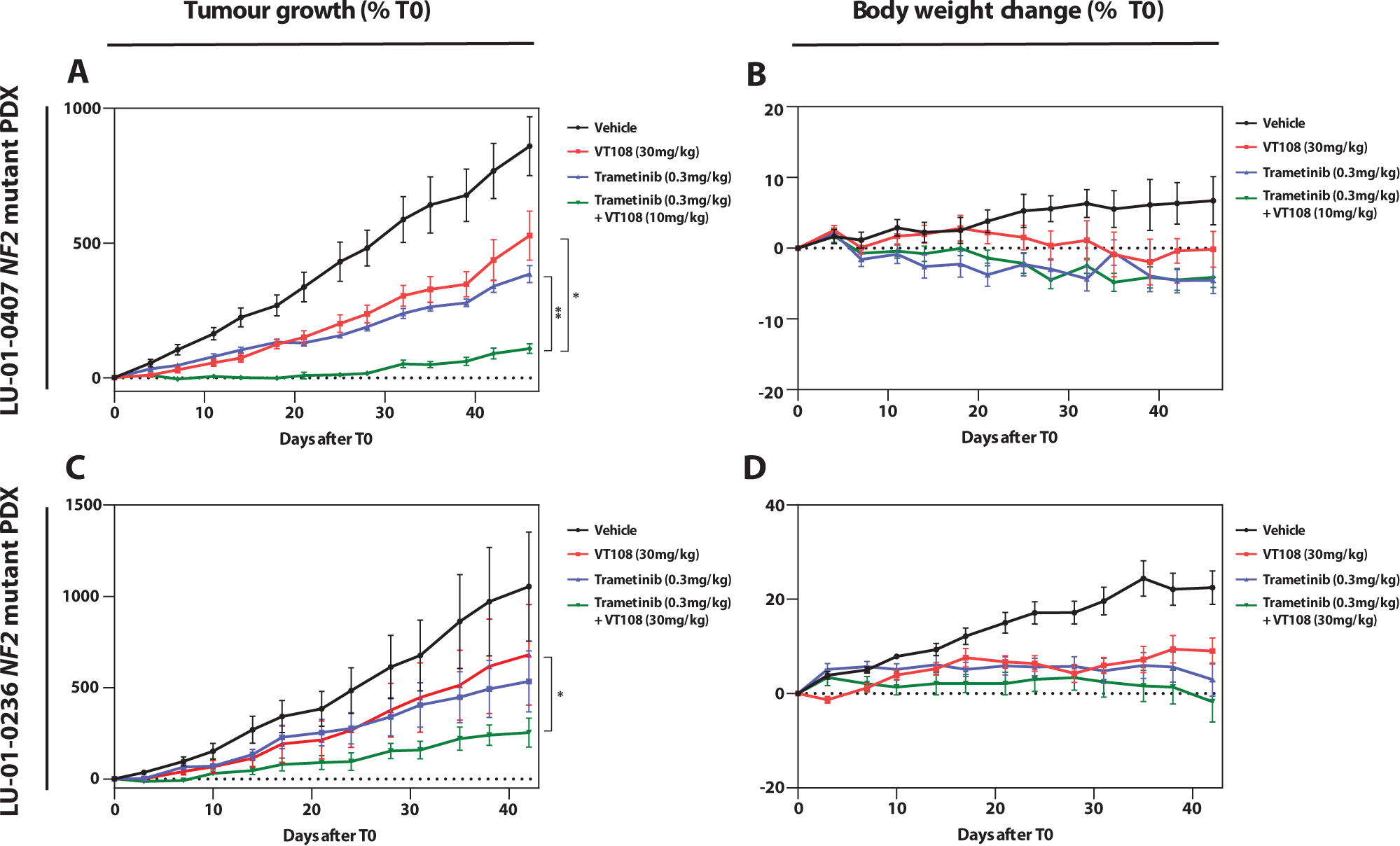
Combined MAPK and Hippo pathway inhibition synergistically impacts NSCLC tumours. **A-D)** The impact of VT108, cobimetinib, or both in combination on Left: the growth of tumours in the NSCLC PDX models LU-01-0407 or LU-01-0236, Right: the average body-weight of each treatment group. n=8 for LU-01-0407, n=6 for LU-01-0236 and error bars represent SEM. The volume of tumours between groups were compared statistically at T42 in a one-way Annova (F-test). * p<0.05, ** p<0.01.

## DISCUSSION

Following its discovery in *Drosophila* as a regulator of tissue growth, the Hippo pathway has been touted as a potential target for new cancer therapies (Dey *et al*., 2020). Two decades later, the first raft of Hippo pathway targeted therapies, which all inhibit YAP/TAZ-TEAD regulated transcription, have entered clinical trials (Calses *et al*., 2019; Pobbati *et al*., 2023). Here, with the aim of identifying potential resistance mechanisms and combination approaches for Hippo targeted therapies, we performed unbiased screens to investigate the cellular response to one such agent, the TEAD palmitoylation inhibitor VT107 (Tang *et al*., 2021). Whole genome CRISR/Cas9 screens of two mesothelioma cell lines with Hippo pathway mutations identified multiple genes that confer either resistance or sensitivity to inhibition of TEAD palmitoylation. Among these were groups of genes that were unique to each cell line and belonged to ontology groups such as fatty acid metabolism (H2052 cells) and interleukin signaling (H226 cells). In addition, select gene ontology groups were commonly enriched in each cell line, including ribosome metabolism and RNA metabolism. Further exploration of these cellular processes is required to understand the molecular mechanisms by which they confer resistance or sensitivity to VT107. On an individual gene basis, we found that a core set of genes modified TEAD’s anti-proliferative effect in both mesothelioma cell lines. Notable among genes that conferred resistance to VT107 was *VGLL4*, which encodes a transcription co-repressor that competes with YAP/TAZ for binding to TEADs. *VGLL4* loss conferred strong resistance to VT107, suggesting that VT107 induces transcription repression by VGLL4/TEAD and associated transcription co-repressors and this is important for VT107’s ability to limit mesothelioma cell proliferation and viability.

Two additional cancer signaling pathways scored strongly in both CRISPR/Cas9 screens: the MAPK and JAK/STAT pathways. The loss of two *bona fide* MAPK pathway tumor suppressors conferred resistance to VT107: *NF1*, which functions in the upstream part of the signal transduction pathway; and *CIC*, a transcription repressor. From the JAK/STAT pathway the upstream signaling repressor SOCS3 conferred VT107 sensitivity, whilst the STAT3 transcription factor conferred resistance to VT107. Interestingly, both STAT3, and, in particular, the MAPK pathway-regulated AP-1 transcription factors, have been reported to co-regulate transcription of many genes (He *et al*, 2021; Koo *et al*., 2020; Obier *et al*., 2016; Pascual *et al*., 2017; Pham *et al*., 2021; Stein *et al*., 2015; Zanconato *et al*., 2015), suggesting a possible mechanism by why which these pathways modulate the cellular impact of VT107. Transcriptome profiling of *NF1* mutant H2052 mesothelioma cells indicated that VT107 strongly represses the expression of many YAP/TEAD target genes in both parental and *NF1* mutant cells, but the transcriptional response is blunted by *NF1* loss. When we examined this further, we found that subsets of genes that are sensitive to YAP/TAZ activity, were partially restored by *NF1* mutation, suggesting that hyperactivation of the MAPK pathway confers resistance to VT107 by reinstating only a subset of YAP/TEAD target genes. The mechanistic basis for this is currently unclear but could potentially be related to the fact that AP-1 transcription factors and YAP/TEAD co-regulate an overlapping transcriptome that is important in cell proliferation and tissue growth (Koo *et al*., 2020; Obier *et al*., 2016; Pascual *et al*., 2017; Pham *et al*., 2021; Stein *et al*., 2015; Zanconato *et al*., 2015). In addition, *NF1* loss could possibly also drive VT107 resistance in H2052 cells by independent mechanisms as it also caused expression changes in many genes that are not regulated by YAP/TAZ-TEAD. The findings of our CRISPR/Cas9 screens and subsequent mechanistic studies spurred us to investigate combination therapies between TEAD and drugs that target either the MAPK or JAK/STAT pathways. Indeed, combining different MEKi’s and TEADi’s synergistically or additively repressed the growth of multiple mesothelioma and NSCLC cell lines and PDX tumors grown in mice, while the JAKi AZD1480 and the TEADi VT107 showed synergistic or strong additive effects on mesothelioma cell lines.

Collectively, our study reports the first genome-wide genetic approaches to investigate the cellular response to chemical inhibition of YAP/TAZ-TEAD-regulated transcription and revealed that modulation of multiple signaling pathways and cellular processes can confer either sensitivity or resistance to VT107. Chief among these is the TEAD binding partner and transcription co-repressor VGLL4, and activity of the MAPK and JAK/STAT pathways. Our experiments provide additional evidence that combining Hippo and MAPK pathway targeted therapies offers promise for the treatment of mesothelioma and other cancers such as NSCLC that have mutations in one or both of these pathways. We also identify the potential of combined targeting of the Hippo and JAK/STAT pathways in mesothelioma, although clearly this requires additional validation. Further, our study identifies potential resistance mechanisms that might limit the effectiveness of TEADi if they fulfil their promise and eventuate as standard of care therapies for one or more cancer types.

## MATERIALS AND METHODS

### Cell culture

NCI-H2052, NCI-H226 and MSTO-211H cells (CVCL_1518, CVCL_1544 and CVCL_1430) were gifted by A/Prof Tom John (Olivia Newton John Cancer Centre, Melbourne, Australia). Cells were cultured in Roswell Park Memorial Institute Medium (Thermo Fisher Scientific) supplemented with 10% Fetal Bovine Serum (Hyclone, SH30396.03) and their mycoplasma-free status was confirmed every 3-6 months with standard PCR (5’ YCGCTGVGTAGTATRYWCGC3’, 5’GCGGTGTGTACAARMCCCGA3’) (Uphoff & Drexler, 2011).

### Proliferation and viability assays

In proliferation assays, cells were seeded in T175 flasks (Cellstar) then treated with DMSO or different doses (within 5 – 2500nM) of VT107 (Vivace Therapeutics) after 24 hours. At 6-days of treatment, the number of cells per condition was counted using a Coulter counter (Beckmann Coulter). In viability assays, cells were seeded in 96-well plates then treated with DMSO or 4 doses of VT107 (11-, 33-, 100-, 300nM) after 24 hours. Cell viability was measured at 96-hours of treatment with an Alamar Blue assay. Measurements were taken at 540/610nm with the Cytation 3 plate reader (Biotek).

### Immunoblotting

Immunoblotting was conducted as in (Zhang *et al*, 2020) and membranes probed with primary antibodies targeting AXL (AB_11217435), CYR61 (AB_2798492), YAP (AB_2218911), p-YAP (AB_2218913), p-ERK (AB_331768), ERK (AB_330744), p-MEK (AB_490903), MEK (AB_10695868), p-STAT3 (AB_2491009), STAT3 (AB_2798995) or NF1 (AB_2798543).

### RNA-sequencing

Cells were treated with 1µM VT107 or the equivalent %(v/v) DMSO and incubated for 24 hours. Total RNA was harvested using Trizol reagent (Thermo Fisher Scientific) and quantified with the Qubit RNA HS (Thermo Fisher Scientific). Indexed libraries were pooled and sequenced on a NextSeq500 (Illumina) and 5-15 million single-end, 75bp reads were generated per sample.

### Analysis of RNA-sequencing data

Differential gene expression (VT107 vs DMSO) was quantified per cell line using the voomwithQualityWeights or EdgeR pipelines on filtered sequencing data (reads with CPM ≥0.5 in all samples) (Liu *et al*, 2015; Ritchie *et al*, 2015). Where comparisons of gene-expression (VT107 vs DMSO) between cell lines were made, differential expression per cell line was quantified using the edgeR pipeline on filtered sequencing data (reads ≥ 10 in at least two biological replicates per group) or an interaction matrix using the limma-voom pipeline was applied. Ranked-list gene set enrichment analysis was conducted with standard protocol (Subramanian *et al*., 2005) using the Hallmark and Oncogenic gene set collections from the Molecular signatures database (MsigDB) (Liberzon *et al*, 2011) or independently selected YAP-TEAD target gene signatures from the literature (Cordenonsi *et al*., 2011; Zanconato *et al*., 2015).

### Proteomics sample preparation

Cells were treated with 1µM VT107 or the equivalent % (v/v) DMSO for 1-hour, 4-hours or 24-hours then lysed by incubation in 4% sodium deoxycholate (Sigma Aldrich) at 95⁰C for 5 minutes. Lysates were tip-probe sonicated (2 x 7 s) at 4⁰C and centrifuged (18,000 x *g* and 4⁰C) for 10 minutes prior to protein quantification by BCA assay (ThermoFisher Scientific). Proteins in lysates (1µg/ml) were reduced by combination with 10mM tris-2-caboxyethyl phosphine (TCEP), then alkylated by incubation at 45°C for 5 minutes in 40mM 2-chloroacetamide (CAA). Reactions were cooled to room temperature then digested by incubation with sequencing-grade Trypsin (Sigma Aldrich) and LysC (Wako, Japan) (1 protease:50 substrate proteins) at 37⁰C for 16 hours. Diluted peptides (50% in isopropanol) were acidified by combining with trifluoroacetic acid (TFA) to a final concentration of 1% (v/v), then loaded directly onto SDB-RPS (Sigma) micro-columns (packed in-house). Loaded columns were washed with isopropanol (supplemented with 1% TFA) followed by 5% acetonitrile (supplemented with 0.2% TFA) prior to elution of peptides in 80% acetonitrile (supplemented with 5% ammonium hydroxide). Eluted peptides were dried by vacuum centrifugation (45°C for 45 min) and resuspended in 2% acetonitrile (supplemented with 0.1% TFA).

### Liquid chromatography tandem mass spectrometry

Peptides were analyzed on a Dionex 3500 nanoHPLC, coupled to an Orbitrap Exploris 480 mass spectrometer (ThermoFisher Scientific) by electrospray ionization in positive mode with 1.9 kV at 275°C. Separation was achieved on a 40 x 75 µm column packed with C18AQ (1.9 µm; Dr Maisch, Germany) at 50s°C for 50 minutes at a flow rate of 300 nL/min. Peptides were eluted over a linear gradient of 3-40% of elution buffer (80% v/v acetonitrile, 0.1% v/v FA). Instruments were operated in data-independent acquisition (DIA) mode with an MS1 spectrum acquired over the mass range 350-950 m/z (60K resolution, 1e6 automatic gain control (AGC) and 50 ms maximum injection time). This was followed by MS/MS analysis of 38 x 16 m/z windows with a 1 m/z overlap via HCD fragmentation (30K resolution, 1e6 AGC) and automatic injection time. Raw MS data were processed using a Sepctronaut DirectDIA (version 14.8.201029.47784) with default parameters and searched against the human UniProt database (October, 2020 release) and filtered to 1% FDR at the peptide spectral match and protein level. The data were searched with a maximum of 2 miss-cleavages, and methionine oxidation and protein N-terminus acetylation were set as variable modifications while carbamidomethylation of cysteine was set as a fixed modification. Quantification was performed using MS2-based extracted ion chromatograms employing 3-6 fragment ions >450 m/z with automated fragment-ion interference removal as described previously (Bruderer *et al*, 2015). Data was analyzed in Perseus and included median normalization and differential expression analysis using t-tests with multiple hypothesis correction using Benjamini-Hochberg FDR adjustment (Tyanova *et al*, 2016).

### Genome-wide CRISPR/Cas9 screens

Two technical replicates were transduced with the genome-wide Brunello CRISPR/Cas9 library at 1000-fold representation and MOI of 0.3 (Doench *et al*., 2016). Positively transduced cells were selected with Puromycin (1µg/ml) for 7 days and maintained at 1000-fold representation for the duration of the screen. Cells were treated with VT107 at its IC_50_ concentration or DMSO (at the equivalent %v/v), for two weeks, then VT107 treatment-concentration increased to its cytostatic dose (and the equivalent %v/v DMSO) for 1-2 weeks. At the treatment endpoints, genomic DNA was extracted from cells at 1000-fold representation using the NucleoSpin Blood XL kit (Clonetech). Sequencing libraries were generated using one-step PCR to amplify the integrated sequence within the construct and the addition of a sample barcode and Illumina adaptors as previously described (Tuano *et al*, 2023). PCR products were purified using AMPure beads (Beckman Coulter) and samples sequenced using HiSeq (Illumina). PoolQ (https://portals.broadinstitute.org/gpp/public/software/poolq) was used for deconvolution of FastQ files and alignment of sgRNA reads. The MAGeCK algorithm was used to identify enriched and depleted genes by comparing sgRNA distribution in drug treated cells to DMSO control (Li *et al*, 2015).

### CRISPR/Cas9 genome editing

The lentiGuide-Puro (Addgene #52963) and lenti-Cas9-2A-Blast (Addgene #73310) vectors were from Addgene. Cells were virally transduced with lenti-Cas9-2A-Blast and selected with Blasticidin-S (3µg/ml) for 7 days. For each gene-of-interest, two single-guide RNAs (sgRNAs) with the highest log_2_fold-change values in the screen were picked from the Brunello CRISPR/Cas9 library (Table 1). Guides were cloned into the lentiGuide-Puro vector as in (Sanjana *et al*, 2014). Cas9-expressing cells were virally transduced with the sgRNA-expression vectors and Puromycin (1µg/ml) selection applied for 7 days. Evidence of CRISPR mutagenesis was obtained through immunoblotting or TIDE analysis of sanger sequencing data at sgRNA-targeted regions (Brinkman & van Steensel, 2019).

### *In vitro* drug efficacy studies

In all studies, cells were incubated at 37°C and 5% CO_2_, plating densities were empirically determined from the doubling rate of individual cell lines and testing compounds applied as dose titrations in matrix format. For combination studies of VT107 (in NCI-H2052 or NCI-H226 cells), 500 cells were seeded per well in 384-well plates and allowed to adhere overnight. The next day, 5 doses of DMSO (within 0.05 – 0.2 %v/v) or 7 doses of VT107 (within 2nM - 5µM) combined with 9 different doses of Trametinib (within 0.6nM-20nM), AZD1480 (within 18.7nM-4nM), Bazedoxifene (within 500-5000nM), BBI-608 (within 25-3500nM), Ruxolitinib (within 1500-50000nM) or Tocilizumab (within 13.8-690nM) were applied to the cells and allowed to incubate with the cells for 6 days. At the incubation endpoint, cells were fixed with 4% paraformaldehyde (Merck life sciences) and stained with 1µg/ml DAPI (Sigma Aldrich). Cell nuclei per condition were quantified using the CellInsight CX7 LED high-content analysis platform according to the manufacturer’s protocol (Thermo Fisher Scientific). Dose-response matrices (%inhibition per treatment) were analyzed using Synergyfinder software (Ianevski *et al*, 2017). For combination treatments of VT108 with cobimetinib (assessed in 42 NSCLC cell lines) or trametinib (assessed in 10 mesothelioma lines), cells were seeded in 384-well plates (NSCLC cells at 250-1000 cells per well; mesothelioma cells at 500-1000 cells per well) and allowed to adhere overnight. The next day (Day 0), 9 doses of VT108 (within 0.05 – 10000nM) in combination with 9 doses of cobimetinib (within 0.4 – 3000nM) or 5 doses of trametinib (within 0.5 – 10000nM) were added to the cells and allowed to incubate with the cells for a total of 5-7 days. Most of the cell lines were incubated with the compounds for 7 days, while a few cell lines that had faster doubling rates were incubated with the compounds for 5 or 6 days. At the end of the assay, cell proliferation was measured by CellTiter-Glo (CTG) Luminescent Cell Viability Assay Kit (Promega) according to the manufacturer’s protocol. The percentage growth inhibition was calculated using Day 0 data as a negative control prior to generating dose response matrices (% inhibition per treatment) for analysis using Synergyfinder.

### *In vivo* drug efficacy studies

All the procedures related to animal handling, care, and the treatment were performed according to the guidelines approved by the Institutional Animal Care and Use Committee (IACUC) of WuXi AppTec following the guidance of the Association for Assessment and Accreditation of Laboratory Animal Care (AAALAC). Trametinib was formulated in vehicle 1 (9% Castor oil+10% PEG-400+81% double distilled water), and VT108 was formulated in vehicle 2 (5% DMSO+10% solutol+85% D5W; D5W=5% glucose). The formulated compounds were orally administered once a day, every day. Animals in the combination groups received the compounds 30 minutes apart – trametinib first, followed by VT108. Animals in the vehicle group received vehicle 1 first and 30 minutes later vehicle 2. In the single agent groups, animals either received formulated trametinib and then vehicle 2, 30 minutes apart or vehicle 1 and then formulated VT108. For the LU-01-0407 study, female BALB/c nude mice were implanted subcutaneously at the right flank with LU-01-0407 tumor slices (∼30 mm^3^) for tumor development. Tumor-bearing animals were randomized, and treatment was started when the average tumor size reached 132 mm^3^. For the LU-01-0236 study, female BALB/c nude mice were implanted subcutaneously at the right flank with the tumor slices (20∼30 mm^3^) for tumor development. Tumor-bearing animals were randomized, and treatment was started when the average tumor size reached 130 mm^3^. Tumor size and animal weights were monitored twice weekly. Tumor volume in mm^3^ was calculated using the formula: V = 0.5 a x b^2^ where a and b are the long and short diameters of the tumor, respectively. Tumor growth inhibition (TGI) in percentage was calculated for each treatment group using the formula: TGI (%) = [1-(Ti-T0)/ (Vi-V0)] ×100, where Ti is the average tumor volume of a treatment group on a given day, T0 is the average tumor volume of the treatment group on the day of treatment start, Vi is the average tumor volume of the vehicle control group on the same day as Ti, and V0 is the average tumor volume of the vehicle group on the day of treatment start. Statistical analysis of difference in the tumor volume among the groups were conducted on the data collected on the indicated treatment day (Day 39 for the LU-01-0407 study; Day 49 for the LU-01-0236 study). A one-way ANOVA was performed to compare the tumor volume among groups, and when a significant F-statistics (a ratio of treatment variance to the error variance) was obtained, comparisons between groups were carried out with Games-Howell test. All data were analyzed using SPSS 17.0. p < 0.05 was considered statistically significant.

## ACKNOWLEDGEMENTS

We thank Avni Anand, Luis Malaver-Ortega, Henry Beetham and members of the Harvey lab for discussions and comments on the manuscript, and A. Chand and M. Shackleton for reagents. K.F.H was supported by a Senior Research Fellowship (APP1078220) and Investigator grant (APP1194467) from the National Health and Medical Research Council of Australia (NHMRC). A.K was partly supported by an Australian Government Research Training Program Scholarship, Rosie Lew Peter MacCallum Cancer Foundation Postgraduate Award. This research was supported by a Lyall Watts Mesothelioma Research Grant from the Cancer Council Victoria (APP1157737) and the Peter MacCallum Cancer Foundation. We acknowledge the Monash Functional Genomics Platform and the following Peter MacCallum Cancer Centre core facilities: Flow Cytometry, Victorian Centre for Functional Genomics, Bioinformatics, Research Laboratory Support Services, and Centre for Advanced Histology and Microscopy, and support to them from the Peter MacCallum Cancer Foundation and the Australian Cancer Research Foundation.

## CONFLICT OF INTEREST

T.T. Tang and L. Post report employment with Vivace Therapeutics and have equity interest in Vivace Therapeutics.

## REFERENCES

1. Brinkman EK, van Steensel B (2019) Rapid Quantitative Evaluation of CRISPR Genome Editing by TIDE and TIDER. Methods Mol Biol 1961: 29–44

2. Bruderer R, Bernhardt OM, Gandhi T, Miladinovic SM, Cheng LY, Messner S, Ehrenberger T, Zanotelli V, Butscheid Y, Escher C et al (2015) Extending the limits of quantitative proteome profiling with data-independent acquisition and application to acetaminophen-treated three-dimensional liver microtissues. Mol Cell Proteomics 14: 1400–1410

3. Bum-Erdene K, Zhou D, Gonzalez-Gutierrez G, Ghozayel MK, Si Y, Xu D, Shannon HE, Bailey BJ, Corson TW, Pollok KE et al (2019) Small-Molecule Covalent Modification of Conserved Cysteine Leads to Allosteric Inhibition of the TEADYap Protein-Protein Interaction. Cell Chem Biol 26: 378–389 e313

4. Calses PC, Crawford JJ, Lill JR, Dey A (2019) Hippo Pathway in Cancer: Aberrant Regulation and Therapeutic Opportunities. Trends Cancer 5: 297–307

5. Chan P, Han X, Zheng B, DeRan M, Yu J, Jarugumilli GK, Deng H, Pan D, Luo X, Wu X (2016) Autopalmitoylation of TEAD proteins regulates transcriptional output of the Hippo pathway. Nat Chem Biol 12: 282–289

6. Cichowski K, Jacks T (2001) NF1 tumor suppressor gene function: narrowing the GAP. Cell 104: 593–604

7. Cordenonsi M, Zanconato F, Azzolin L, Forcato M, Rosato A, Frasson C, Inui M, Montagner M, Parenti AR, Poletti A et al (2011) The Hippo transducer TAZ confers cancer stem cell-related traits on breast cancer cells. Cell 147: 759–772

8. Davis JR, Tapon N (2019) Hippo signalling during development. Development 146

9. Dey A, Varelas X, Guan KL (2020) Targeting the Hippo pathway in cancer, fibrosis, wound healing and regenerative medicine. Nat Rev Drug Discov 19: 480–494

10. Doench JG, Fusi N, Sullender M, Hegde M, Vaimberg EW, Donovan KF, Smith I, Tothova Z, Wilen C, Orchard R et al (2016) Optimized sgRNA design to maximize activity and minimize off-target effects of CRISPR-Cas9. Nat Biotechnol 34: 184–191

11. Doench JG, Hartenian E, Graham DB, Tothova Z, Hegde M, Smith I, Sullender M, Ebert BL, Xavier RJ, Root DE (2014) Rational design of highly active sgRNAs for CRISPR-Cas9-mediated gene inactivation. Nat Biotechnol 32: 1262–1267

12. Gibault F, Sturbaut M, Coevoet M, Pugniere M, Burtscher A, Allemand F, Melnyk P, Hong W, Rubin BP, Pobbati AV et al (2021) Design, Synthesis and Evaluation of a Series of 1,5-Diaryl-1,2,3-triazole-4-carbohydrazones as Inhibitors of the YAP-TAZ/TEAD Complex. ChemMedChem 16: 2823–2844

13. Gridnev A, Maity S, Misra JR (2022) Structure-based discovery of a novel small-molecule inhibitor of TEAD palmitoylation with anticancer activity. Front Oncol 12: 1021823

14. Halder G, Johnson RL (2011) Hippo signaling: growth control and beyond. Development 138: 9–22

15. Hao Y, Chun A, Cheung K, Rashidi B, Yang X (2008) Tumor suppressor LATS1 is a negative regulator of oncogene YAP. J Biol Chem 283: 5496–5509

16. Hart T, Chandrashekhar M, Aregger M, Steinhart Z, Brown KR, MacLeod G, Mis M, Zimmermann M, Fradet-Turcotte A, Sun S et al (2015) High-Resolution CRISPR Screens Reveal Fitness Genes and Genotype-Specific Cancer Liabilities. Cell 163: 1515–1526

17. Harvey KF, Hariharan IK (2012) The hippo pathway. Cold Spring Harb Perspect Biol 4: a011288

18. Harvey KF, Zhang X, Thomas DM (2013) The Hippo pathway and human cancer. Nat Rev Cancer 13: 246–257

19. He L, Pratt H, Gao M, Wei F, Weng Z, Struhl K (2021) YAP and TAZ are transcriptional co-activators of AP-1 proteins and STAT3 during breast cellular transformation. Elife 10

20. Heinrich PC, Behrmann I, Muller-Newen G, Schaper F, Graeve L (1998) Interleukin-6-type cytokine signalling through the gp130/Jak/STAT pathway. Biochem J 334 (Pt 2): 297–314

21. Herranz H, Hong X, Cohen SM (2012) Mutual repression by bantam miRNA and Capicua links the EGFR/MAPK and Hippo pathways in growth control. Current Biology 22: 651–657

22. Holden JK, Crawford JJ, Noland CL, Schmidt S, Zbieg JR, Lacap JA, Zang R, Miller GM, Zhang Y, Beroza P et al (2020) Small Molecule Dysregulation of TEAD Lipidation Induces a Dominant-Negative Inhibition of Hippo Pathway Signaling. Cell Rep 31: 107809

23. Hong JH, Hwang ES, McManus MT, Amsterdam A, Tian Y, Kalmukova R, Mueller E, Benjamin T, Spiegelman BM, Sharp PA et al (2005) TAZ, a transcriptional modulator of mesenchymal stem cell differentiation. Science 309: 1074–1078

24. Hu L, Sun Y, Liu S, Erb H, Singh A, Mao J, Luo X, Wu X (2022) Discovery of a new class of reversible TEA domain transcription factor inhibitors with a novel binding mode. Elife 11

25. Ianevski A, He L, Aittokallio T, Tang J (2017) SynergyFinder: a web application for analyzing drug combination dose-response matrix data. Bioinformatics 33: 2413–2415

26. Kaneda A, Seike T, Danjo T, Nakajima T, Otsubo N, Yamaguchi D, Tsuji Y, Hamaguchi K, Yasunaga M, Nishiya Y et al (2020) The novel potent TEAD inhibitor, K-975, inhibits YAP1/TAZ-TEAD protein-protein interactions and exerts an anti-tumor effect on malignant pleural mesothelioma. Am J Cancer Res 10: 4399–4415

27. Kawamura-Saito M, Yamazaki Y, Kaneko K, Kawaguchi N, Kanda H, Mukai H, Gotoh T, Motoi T, Fukayama M, Aburatani H et al (2006) Fusion between CIC and DUX4 up-regulates PEA3 family genes in Ewing-like sarcomas with t(4;19)(q35;q13) translocation. Hum Mol Genet 15: 2125–2137

28. Kim JW, Ponce RK, Okimoto RA (2021) Capicua in Human Cancer. Trends Cancer 7: 77–86

29. Kim MH, Kim J, Hong H, Lee SH, Lee JK, Jung E, Kim J (2016) Actin remodeling confers BRAF inhibitor resistance to melanoma cells through YAP/TAZ activation. The EMBO journal 35: 462–478

30. Koo JH, Guan KL (2018) Interplay between YAP/TAZ and Metabolism. Cell Metab 28: 196–206

31. Koo JH, Plouffe SW, Meng Z, Lee DH, Yang D, Lim DS, Wang CY, Guan KL (2020) Induction of AP-1 by YAP/TAZ contributes to cell proliferation and organ growth. Genes Dev 34: 72–86

32. Koontz LM, Liu-Chittenden Y, Yin F, Zheng Y, Yu J, Huang B, Chen Q, Wu S, Pan D (2013) The Hippo effector Yorkie controls normal tissue growth by antagonizing scalloped-mediated default repression. Dev Cell 25: 388–401

33. Kulkarni A, Chang MT, Vissers JHA, Dey A, Harvey KF (2020) The Hippo Pathway as a Driver of Select Human Cancers. Trends Cancer

34. Laraba L, Hillson L, de Guibert JG, Hewitt A, Jaques MR, Tang TT, Post L, Ercolano E, Rai G, Yang SM et al (2023) Inhibition of YAP/TAZ-driven TEAD activity prevents growth of NF2-null schwannoma and meningioma. Brain 146: 1697–1713

35. LeBlanc VG, Firme M, Song J, Chan SY, Lee MH, Yip S, Chittaranjan S, Marra MA (2017) Comparative transcriptome analysis of isogenic cell line models and primary cancers links capicua (CIC) loss to activation of the MAPK signalling cascade. J Pathol 242: 206–220

36. Li Q, Sun Y, Jarugumilli GK, Liu S, Dang K, Cotton JL, Xiol J, Chan PY, DeRan M, Ma L et al (2020) Lats1/2 Sustain Intestinal Stem Cells and Wnt Activation through TEAD-Dependent and Independent Transcription. Cell Stem Cell 26: 675–692 e678

37. Li W, Koster J, Xu H, Chen CH, Xiao T, Liu JS, Brown M, Liu XS (2015) Quality control, modeling, and visualization of CRISPR screens with MAGeCK-VISPR. Genome Biol 16: 281

38. Li W, Xu H, Xiao T, Cong L, Love MI, Zhang F, Irizarry RA, Liu JS, Brown M, Liu XS (2014) MAGeCK enables robust identification of essential genes from genome-scale CRISPR/Cas9 knockout screens. Genome Biol 15: 554

39. Liberzon A, Subramanian A, Pinchback R, Thorvaldsdottir H, Tamayo P, Mesirov JP (2011) Molecular signatures database (MSigDB) 3.0. Bioinformatics 27: 1739–1740

40. Lin L, Sabnis AJ, Chan E, Olivas V, Cade L, Pazarentzos E, Asthana S, Neel D, Yan JJ, Lu X (2015) The Hippo effector YAP promotes resistance to RAF-and MEK-targeted cancer therapies. Nature genetics 47: 250–256

41. Liu R, Holik AZ, Su S, Jansz N, Chen K, Leong HS, Blewitt ME, Asselin-Labat ML, Smyth GK, Ritchie ME (2015) Why weight? Modelling sample and observational level variability improves power in RNA-seq analyses. Nucleic Acids Res 43: e97

42. Lu T, Li Y, Lu W, Spitters T, Fang X, Wang J, Cai S, Gao J, Zhou Y, Duan Z et al (2021) Discovery of a subtype-selective, covalent inhibitor against palmitoylation pocket of TEAD3. Acta Pharm Sin B 11: 3206–3219

43. Lu W, Fan M, Ji W, Tse J, You I, Ficarro SB, Tavares I, Che J, Kim AY, Zhu X et al (2023) Structure-Based Design of Y-Shaped Covalent TEAD Inhibitors. J Med Chem 66: 4617–4632

44. Lu W, Wang J, Li Y, Tao H, Xiong H, Lian F, Gao J, Ma H, Lu T, Zhang D et al (2019) Discovery and biological evaluation of vinylsulfonamide derivatives as highly potent, covalent TEAD autopalmitoylation inhibitors. Eur J Med Chem 184: 111767

45. Luscan A, Shackleford G, Masliah-Planchon J, Laurendeau I, Ortonne N, Varin J, Lallemand F, Leroy K, Dumaine V, Hivelin M et al (2014) The activation of the WNT signaling pathway is a Hallmark in neurofibromatosis type 1 tumorigenesis. Clin Cancer Res 20: 358–371

46. Manning SA, Kroeger B, Harvey KF (2020) The regulation of Yorkie, YAP and TAZ: new insights into the Hippo pathway. Development 147

47. Miyanaga A, Masuda M, Tsuta K, Kawasaki K, Nakamura Y, Sakuma T, Asamura H, Gemma A, Yamada T (2015) Hippo pathway gene mutations in malignant mesothelioma: revealed by RNA and targeted exon sequencing. J Thorac Oncol 10: 844–851

48. Murakami H, Mizuno T, Taniguchi T, Fujii M, Ishiguro F, Fukui T, Akatsuka S, Horio Y, Hida T, Kondo Y et al (2011) LATS2 is a tumor suppressor gene of malignant mesothelioma. Cancer Res 71: 873–883

49. Noland CL, Gierke S, Schnier PD, Murray J, Sandoval WN, Sagolla M, Dey A, Hannoush RN, Fairbrother WJ, Cunningham CN (2016) Palmitoylation of TEAD Transcription Factors Is Required for Their Stability and Function in Hippo Pathway Signaling. Structure 24: 179–186

50. Obier N, Cauchy P, Assi SA, Gilmour J, Lie ALM, Lichtinger M, Hoogenkamp M, Noailles L, Cockerill PN, Lacaud G et al (2016) Cooperative binding of AP-1 and TEAD4 modulates the balance between vascular smooth muscle and hemogenic cell fate. Development 143: 4324–4340

51. Park J, Eisenbarth D, Choi W, Kim H, Choi C, Lee D, Lim D-S (2020) YAP and AP-1 Cooperate to Initiate Pancreatic Cancer Development from Ductal Cells in Mice. Cancer Research 80: 4768–4779

52. Pascual J, Jacobs J, Sansores-Garcia L, Natarajan M, Zeitlinger J, Aerts S, Halder G, Hamaratoglu F (2017) Hippo reprograms the transcriptional response to Ras signaling. Developmental cell 42: 667–680. e664

53. Pham TH, Hagenbeek TJ, Lee HJ, Li J, Rose CM, Lin E, Yu M, Martin SE, Piskol R, Lacap JA et al (2021) Machine-Learning and Chemicogenomics Approach Defines and Predicts Cross-Talk of Hippo and MAPK Pathways. Cancer Discov 11: 778–793

54. Pobbati AV, Han X, Hung AW, Weiguang S, Huda N, Chen GY, Kang C, Chia CS, Luo X, Hong W et al (2015) Targeting the Central Pocket in Human Transcription Factor TEAD as a Potential Cancer Therapeutic Strategy. Structure 23: 2076–2086

55. Pobbati AV, Kumar R, Rubin BP, Hong W (2023) Therapeutic targeting of TEAD transcription factors in cancer. Trends Biochem Sci 48: 450–462

56. Ratner N, Miller SJ (2015) A RASopathy gene commonly mutated in cancer: the neurofibromatosis type 1 tumour suppressor. Nat Rev Cancer 15: 290–301

57. Rawlings JS, Rosler KM, Harrison DA (2004) The JAK/STAT signaling pathway. Journal of cell science 117: 1281–1283

58. Reddy BV, Irvine KD (2013) Regulation of Hippo signaling by EGFR-MAPK signaling through Ajuba family proteins. Dev Cell 24: 459–471

59. Rice KD, Aay N, Anand NK, Blazey CM, Bowles OJ, Bussenius J, Costanzo S, Curtis JK, Defina SC, Dubenko L et al (2012) Novel Carboxamide-Based Allosteric MEK Inhibitors: Discovery and Optimization Efforts toward XL518 (GDC-0973). ACS Med Chem Lett 3: 416–421

60. Ritchie ME, Phipson B, Wu D, Hu Y, Law CW, Shi W, Smyth GK (2015) limma powers differential expression analyses for RNA-sequencing and microarray studies. Nucleic Acids Res 43: e47

61. Sanjana NE, Shalem O, Zhang F (2014) Improved vectors and genome-wide libraries for CRISPR screening. Nat Methods 11: 783–784

62. Sekido Y, Pass HI, Bader S, Mew DJ, Christman MF, Gazdar AF, Minna JD (1995) Neurofibromatosis type 2 (NF2) gene is somatically mutated in mesothelioma but not in lung cancer. Cancer Res 55: 1227–1231

63. Stein C, Bardet AF, Roma G, Bergling S, Clay I, Ruchti A, Agarinis C, Schmelzle T, Bouwmeester T, Schubeler D et al (2015) YAP1 Exerts Its Transcriptional Control via TEAD-Mediated Activation of Enhancers. PLoS genetics 11: e1005465

64. Subramanian A, Tamayo P, Mootha VK, Mukherjee S, Ebert BL, Gillette MA, Paulovich A, Pomeroy SL, Golub TR, Lander ES et al (2005) Gene set enrichment analysis: a knowledge-based approach for interpreting genome-wide expression profiles. Proc Natl Acad Sci U S A 102: 15545–15550

65. Sun Y, Hu L, Tao Z, Jarugumilli GK, Erb H, Singh A, Li Q, Cotton JL, Greninger P, Egan RK et al (2022) Pharmacological blockade of TEAD-YAP reveals its therapeutic limitation in cancer cells. Nat Commun 13: 6744

66. Tang TT, Konradi AW, Feng Y, Peng X, Ma M, Li J, Yu FX, Guan KL, Post L (2021) Small Molecule Inhibitors of TEAD Auto-palmitoylation Selectively Inhibit Proliferation and Tumor Growth of NF2-deficient Mesothelioma. Mol Cancer Ther

67. Tapon N, Harvey KF, Bell DW, Wahrer DC, Schiripo TA, Haber DA, Hariharan IK (2002) salvador Promotes both cell cycle exit and apoptosis in Drosophila and is mutated in human cancer cell lines. Cell 110: 467–478

68. Tuano NK, Beesley J, Manning M, Shi W, Perlaza-Jimenez L, Malaver-Ortega LF, Paynter JM, Black D, Civitarese A, McCue K et al (2023) CRISPR screens identify gene targets at breast cancer risk loci. Genome Biol 24: 59

69. Tyanova S, Temu T, Sinitcyn P, Carlson A, Hein MY, Geiger T, Mann M, Cox J (2016) The Perseus computational platform for comprehensive analysis of (prote)omics data. Nat Methods 13: 731–740

70. Uphoff CC, Drexler HG (2011) Detecting mycoplasma contamination in cell cultures by polymerase chain reaction. Methods Mol Biol 731: 93–103

71. Varelas X, Sakuma R, Samavarchi-Tehrani P, Peerani R, Rao BM, Dembowy J, Yaffe MB, Zandstra PW, Wrana JL (2008) TAZ controls Smad nucleocytoplasmic shuttling and regulates human embryonic stem-cell self-renewal. Nat Cell Biol 10: 837–848

72. Wang W, Huang J, Wang X, Yuan J, Li X, Feng L, Park JI, Chen J (2012) PTPN14 is required for the density-dependent control of YAP1. Genes Dev 26: 1959–1971

73. Wang Y, Xu X, Maglic D, Dill MT, Mojumdar K, Ng PK, Jeong KJ, Tsang YH, Moreno D, Bhavana VH et al (2018) Comprehensive Molecular Characterization of the Hippo Signaling Pathway in Cancer. Cell Rep 25: 1304–1317 e1305

74. Xu GF, O’Connell P, Viskochil D, Cawthon R, Robertson M, Culver M, Dunn D, Stevens J, Gesteland R, White R et al (1990) The neurofibromatosis type 1 gene encodes a protein related to GAP. Cell 62: 599–608

75. Zanconato F, Forcato M, Battilana G, Azzolin L, Quaranta E, Bodega B, Rosato A, Bicciato S, Cordenonsi M, Piccolo S (2015) Genome-wide association between YAP/TAZ/TEAD and AP-1 at enhancers drives oncogenic growth. Nat Cell Biol 17: 1218–1227

76. Zhang X, Yang L, Szeto P, Abali GK, Zhang Y, Kulkarni A, Amarasinghe K, Li J, Vergara IA, Molania R et al (2020) The Hippo pathway oncoprotein YAP promotes melanoma cell invasion and spontaneous metastasis. Oncogene

77. Zhao B, Ye X, Yu J, Li L, Li W, Li S, Yu J, Lin JD, Wang CY, Chinnaiyan AM et al (2008) TEAD mediates YAP-dependent gene induction and growth control. Genes Dev 22: 1962–1971

78. Zheng Y, Pan D (2019) The Hippo Signaling Pathway in Development and Disease. Dev Cell 50: 264–282

79. Zheng ZY, Anurag M, Lei JT, Cao J, Singh P, Peng J, Kennedy H, Nguyen NC, Chen Y, Lavere P et al (2020) Neurofibromin Is an Estrogen Receptor-alpha Transcriptional Co-repressor in Breast Cancer. Cancer Cell 37: 387–402 e387

